# Reversion of the photoperiod dependence of flowering in rice with synthetic Hd1-microProteins

**DOI:** 10.1101/266486

**Authors:** Tenai Eguen, Jorge Gomez Ariza, Kaushal Kumar Bhati, Bin Sun, Fabio Fornara, Stephan Wenkel

## Abstract

Rice (*Oryza sativa*) is a facultative short-day plant that flowers very late when grown in non-inductive long day conditions. Photoperiod-dependent flowering in rice is regulated by heading date (Hd1) which acts as both an activator and repressor of flowering in a day length-dependent manner. In order to regulate flowering of rice in long days (LD), overexpression of a synthetic Hd1miP, which is capable of interacting with Hd1, was employed. Transgenic *Hd1miP* rice plants flowered significantly earlier when grown in LD compared to SD, showing that synthetic microProteins can be used to revert short-day plants to long-day plants. Yield analysis revealed that although the *OX-Hd1miP* grains are comparable to WT in terms of the size of the grains produced, *OX-Hd1miP* plants like *hd1* knockout plants, are compromised in the number of grains produced and the grain maturity rate, suggesting an additional unrecognized role of Hd1 in grain maturity.

## INTRODUCTION

The timing of the transition from vegetative to reproductive growth in plants is essential to its reproductive success (GARNER & ALLARD, 1920; Yanovsky & Kay, 2003). The transition to the reproductive state is dependent on both external and internal factors, one important factor being the length of the day (photoperiod). In the model organism *Arabidopsis thaliana*, B-Box zinc finger transcription factor CONSTANS (CO) positively regulates FLOWERING LOCUS-T (FT) to activate flowering in response to long days (Kardailsky et al., 1999; Putterill, Robson, Lee, Simon, & Coupland, 1995; Suárez-López et al., 2001; Valverde, 2011):

Flowering in rice, a facultative short day plant is modulated by HEADING DATE 1 (Hd1). Hd1 is an ortholog of CO and although both proteins regulate flowering through a similar mechanism, Hd1 regulation of flowering features some unique differences. Flowering control in rice consists of two major connected pathways, Hd1 mediated and EARLY HEADING DATE 1 (Ehd1) mediated pathways (Doi et al., 2004; Izawa et al., 2002; Komiya, Ikegami, Tamaki, Yokoi, & Shimamoto, 2008). In SD conditions, Hd1 functions as a flowering activator and in conjunction with Ehd1 acts to positively regulate *RICE FLOWERING LOCUS T 1* (RFT1) and Heading date 3a (Hd3a), the latter of which are florigens important for rice transition to the reproductive phase (Corbesier et al., 2007; Doi et al., 2004; Hayama, Yokoi, Tamaki, Yano, & Shimamoto, 2003; Kojima et al., 2002; Komiya et al., 2008; Tamaki, Matsuo, Wong, Yokoi, & Shimamoto, 2007; Yano et al., 2000). In long day (LD) conditions however, Hd1 unlike CO, switches to a floral repressor thereby inhibiting Ehd1 dependent pathway and Hd3a transcription; this results in a significantly late flowering phenotype (Hayama et al., 2003; R. Ishikawa et al., 2011; Ryo Ishikawa et al., 2005; Izawa et al., 2002; Nemoto, Nonoue, Yano, & Izawa, 2016). Hd1’s role in flowering is clearly observed in knockout and overexpressor plants of *Hd1* which show early and late flowering phenotype respectively in comparison to WT in LD conditions (Gómez-Ariza et al., 2015; R. Ishikawa et al., 2011; Izawa et al., 2002; Takahashi, Teshima, Yokoi, Innan, & Shimamoto, 2009; Wang et al., 2013; Yano et al., 2000; Zhang et al., 2016).

Another difference between Hd1 and CO is the presence of microProtein-mediated inhibition of flowering in Arabidopsis in LD conditions. MicroProteins (miPs) are small single domain proteins that interfere with larger multi-domain proteins to regulate their activity by inactivating or altering their target’s function (Eguen, Straub, Graeff, & Wenkel, 2015; M. Graeff & Wenkel, 2012; Staudt & Wenkel, 2011). In Arabidopsis two microProteins were recently discovered as regulators of flowering, microProtein 1a (miP1a) and microProtein1b (miP1b). MiP1a and miP1b are homologous B-Box proteins that regulate flowering by binding to and inhibiting the function of larger transcription factor CO, a positive regulator of flowering. MiP1a/b inhibition requires the activity of the repressor TOPLESS (Moritz Graeff et al., 2016).

Attempts to identify miP1a/b orthologs in rice revealed that Hd1-related miPs do not exist in monocotyledonous plants. Previous studies in Arabidopsis however, show that the overexpression of a truncated splice variant of CO, which contains the B-Box domain but lacks both the middle low complexity region and the CCT domain can mimic the miP1a/b and *CO* knockout by exhibiting late flowering phenotype in LD conditions. This suggests that the B-box only splice variant of CO may inhibit the activity of the full length CO in a microProtein-like manner (Moritz Graeff et al., 2016).

In order to control HD1 flowering in long day we created a synthetic Hd1 microProtein in *O. sativa*. In this study we show that the synthetic Hd1 microProtein (Hd1miP) is capable of interacting with Hd1 and when overexpressed in rice inhibits the repressor function of Hd1 in LD conditions, while having comparable flowering phenotype to WT in SD. Furthermore analysis of the yield physical characteristics of *Hd1miP* overexpressors, *Hd1* knockout and *Hd1* overexpressors reveals that although the grain maturity rate and fertility of OX-*Hd1miP* plants are slightly compromised, the other characteristics such as the number of panicles produced and grain size are similar to WT suggesting a possible role of Hd1 in grain maturity rate and fertility.

## RESULTS AND DISCUSSION

### Overexpression of Hd1 inhibits late flowering phenotype in long day

The blast analysis of Hd1 reveals that there are no B-Box related miPs in *O. sativa* (data not shown). In order to determine the effects of a floral synthetic miP in rice, the full and truncated forms of the *Hd1* (LOC_Os06g16370) was expressed under the constitutive rice ACTIN2 promoter. The truncated form (*Hd1miP*) consists of a 3’ *3xFLAG* coding sequence (CDS) and the CDS of *Hd1* from 4- 429bp downstream of the start; this codes for the two Hd1 B-Box domains fused to an N-terminal FLAG tag but lacks the middle region and the CCT-domain. As a control the full length *Hd1* was also overexpressed under the ACTIN2 promoter and consists of a 3x*FLAG* CDS and the entire *HD1* CDS (Figure 1a). The protein expression of three *Hd1miP* overexpressors and three full-length *Hd1* overexpressors were measured and the highest overexpressor of Hd1 (*OX-Hd1*) and the 2 highest overexpressors of Hd1miP (*OX-Hd1miP-1 and OX-Hd1miP-2*) were selected for further experiments (Figure 1b).

**Figure 1:**
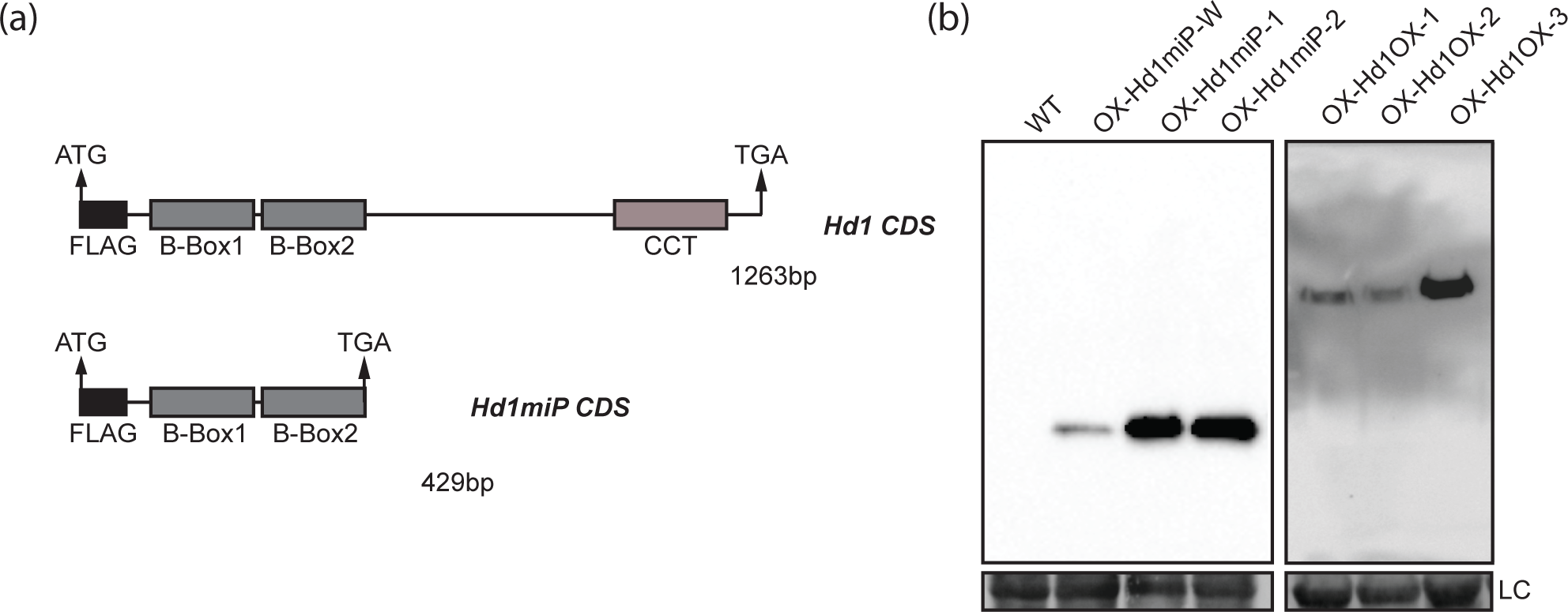
The construction of Hd1miP. (A) Schematic diagram of Hd1 and Hd1miP including the FLAG, B-Box and CCT domain. (B) The protein expression profile of Hd1 and Hd1miP overexpression lines. LC-loading control.

The flowering phenotype of *Hd1* overexpressors and *Hd1* knockout plants has been previously described (R. Ishikawa et al., 2011; Izawa et al., 2002; Takahashi et al., 2009; Yano et al., 2000). To determine the effect of overexpression *Hd1miP* in flowering, we measured the flowering time of WT, *OX-Hd1miP*, *OX-Hd1* and *hd1-1* in SD (10L), LD (14.5L) and strong long day (SLD-16.5L). We observed that in SD there is no significant difference between WT, *OX-Hd1miP* and *OX-Hd1, hd1-1*. In LD, we observed that as expected, *hd1-1* flowered much earlier than WT. *OX-Hd1miP* also flowered earlier than WT while *OX-Hd1* did not show significant differences compared to WT; both *OX-Hd1* and WT flowered late in LD conditions. In SLD, *OX-Hd1miP* and *hd1-1* showed significantly earlier flowering than WT. *OX-Hd1* on the other hand, showed significantly later flowering phenotype than WT (Figure 2). The flowering phenotypes reveal a possible mechanism by which Hd1miP can inhibit the long day phenotype in the presence of a WT Hd1. This mechanism is miP-like in nature as it reveals the post-translation regulation of a related endogenous target via compatible domain interaction, the output of which can be observed in Hd1miPs ability to significantly affect flowering in LD. Although the effect of the Hd1 levels can be seen in LD conditions they were not seen in the short day where all the lines showed similar flowering time this suggests that Hd1’s function may primarily be as a repressor of flowering in LD. Recent studies have also shown the power of synthetic miPs in regulating different processes in Arabidopsis (Dolde et al., 2018).

**Figure 2:**
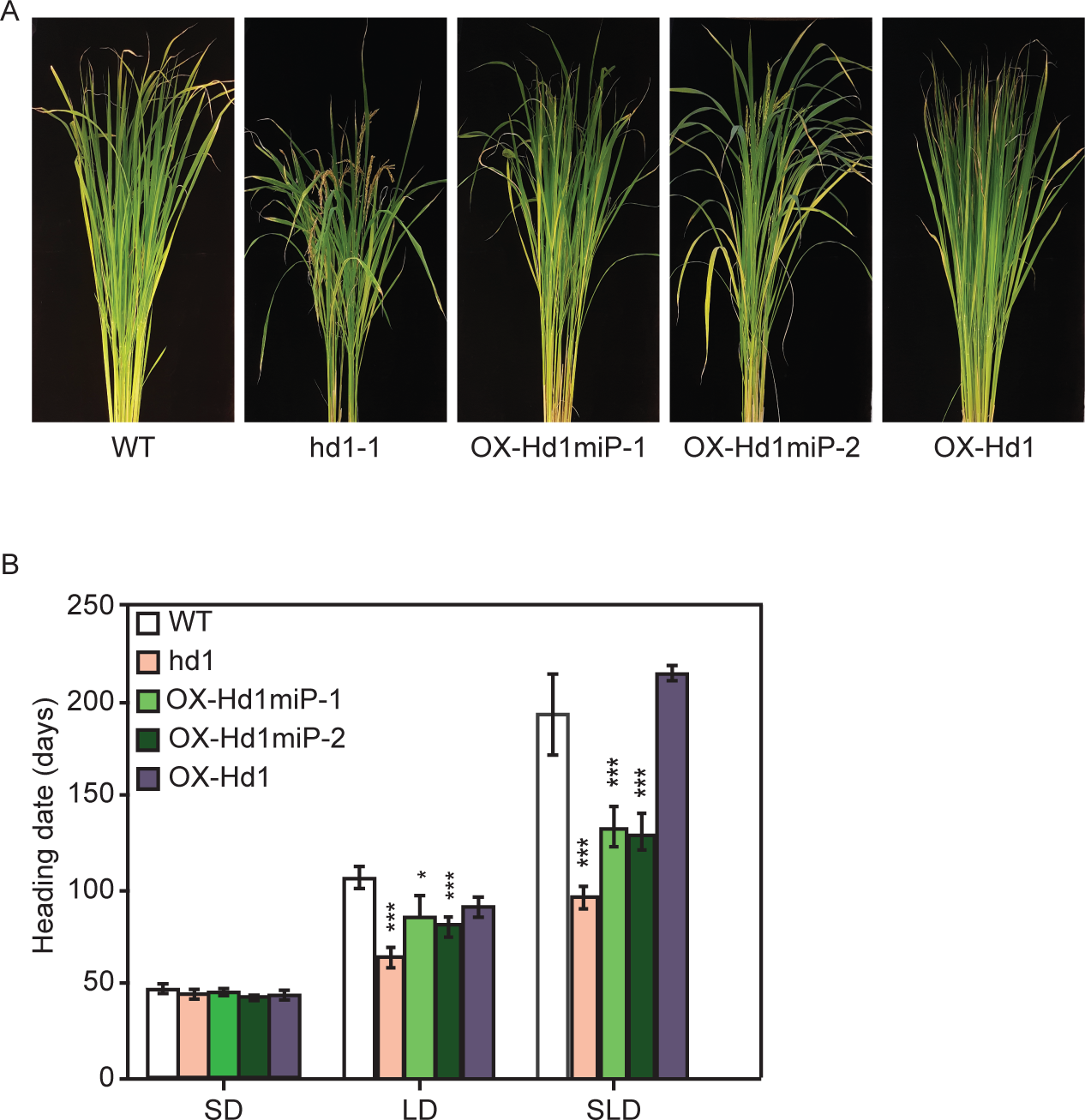
Flowering phenotype of Hd1miP overexpressors (A) Picture of WT, hd1-1, OX-Hd1miP, OX-Hd1 in rice plants grown in LD conditions. (B) Heading date count of WT, hd1-1, OX-Hd1miP, OX-Hd1 in Short day-SD (10L), long day –LD (14.5L) and strong long day SLD (16.5L)

### Hd1miP interacts with Hd1 full length protein

B-Box proteins are known to interact with other B-Box proteins via the B-Box domain (Gangappa & Botto, 2014). In order to determine if the Hd1miP is capable of interacting with Hd1, their interaction was tested in yeast. Hd1miP was fused to the GAL4 binding domain (BD) while the full length Hd1 was fused to the GAL4 activation domain (AD). We observed that Hd1miP was able to interact with Hd1 full length both on non-selective SD-medium lacking leucine and tryptophan (-L/-W), and on media lacking leucine, histidine and tryptophan (-L/-H/-W) in the presence of 10mM 3-aminotriazole. Negative control lack-of-interaction analysis of the Hd1miP with the AD-vector and Hd1 with the BD-vector shows Hd1- Hd1miP interaction to be specific (Figure 3). This further reinforces the idea that Hd1miP regulation of flowering is through a post-translational miP-like inhibition of Hd1 function.

**Figure 3:**
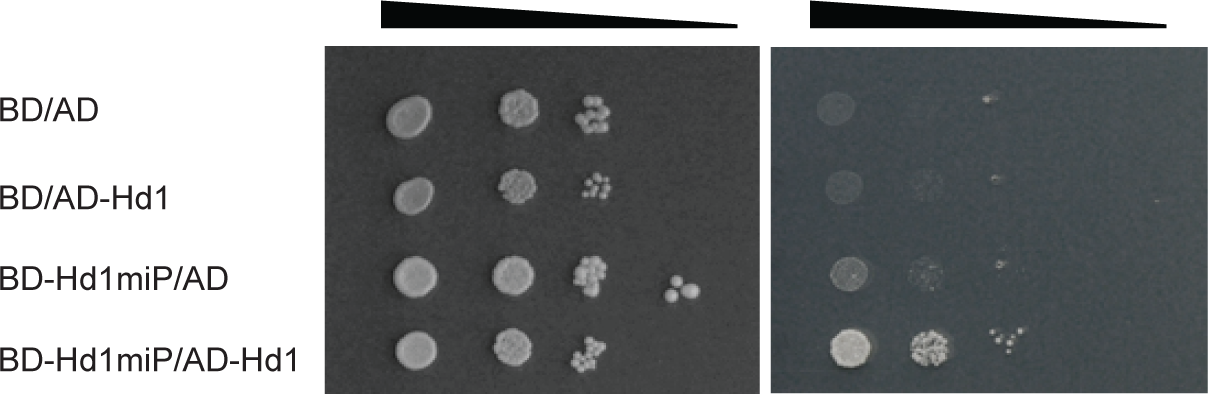
Interaction of Hd1miP and Hd1 proteins in vivo. Interaction of Hd1miP and Hd1 was tested using a yeast two-hybrid assay. Yeast growth was observed in serial dilutions on non-selective media lacking leucine and tryptophan (SD-L-W). Positive interactors showed growth on selective media lacking leucine, tryptophan, and histidine (SD-L-W-H) supplemented with 10mM 3-aminotriazole (3-AT).

### Hd1 levels affect grain and maturity but not grain size

Hd1’s role in development is not limited to flowering but also rice yield and performance (Zhang et al., 2016). To determine the viability and productivity of *OX-Hd1miP* plants, the grain yield of *OX-Hd1miP* plants was compared to WT, *OX-Hd1* and *hd1-1* with regard to number of panicles/plant, number of grains per plant, fertility rate, moisture content and length and width of grains. The measurements were obtained 35 days after the first panicle emergence of each plant grown in LD conditions. The number of panicles produced by *OX-Hd1miP*, *OX-Hd1*, *hd1-1* and WT showed no significant difference indicating that Hd1 levels/activity does not affect the number of panicles produced (Table 1).

The panicles produced were further classified based on the maturity of the panicles. The maturity was visually determined based on the following criteria. Panicles that were greater than 90% green were considered as immature. Panicles that were 70-90% green were considered semi-mature while panicles that were yellowish-brown were considered as mature. Over 50% of the WT panicles were mature in WT plants at the 35 days harvesting time point. In *hd1-1* plants, an average of 36% of the panicles was mature, showing a significantly slower maturity rate in *hd1-1* plants. In *Hd1miP-1* plants an average of 33% of the panicles were mature at harvest, while in *Hd1miP-2* 38% of the panicles were mature which is also less than WT. *OX-Hd1* on the other hand exhibited an average of about 70% mature panicles suggesting that Hd1 may either play a direct or indirect role in the maturity rate of the panicles. Additionally, extending the harvest time beyond 35 days after first panicle emergence may significantly increase the yield of *OX-Hd1miP* plants (Table 1).

Analysis of the fertility rate of the different plants was measured by calculating the ratio of the filled grains to unfilled grains per mature panicle. *OX-Hd1miP* plants and *hd1-1* plants showed slightly lower but significant fertility rate compared to WT, while *OX-Hd1* fertility rate was comparable to WT. Additionally the average total number of grains produced in all the lines were less than that of WT plants. This shows that the levels/activity of Hd1 also affects the fertility rate and the number of grains produced in LD. This regulation occurs irrespective of if Hd1 is inhibited at a transcriptional (*hd1-1*) or at a post translational level (*OX-Hd1miP*) (Table 1).

Hd1 levels/activity also seems to affect the moisture content of the Hd1 grains. The knockout mutant *hd1-1* had the highest moisture content. *OX-Hd1miP* had slightly higher but significant moisture content, while *OX-Hd1* had comparable moisture level compared to WT plants. (Figure 4a) It is feasible that the delayed maturity of the grains could be attributed to the moisture content of the grains.

**Figure 4:**
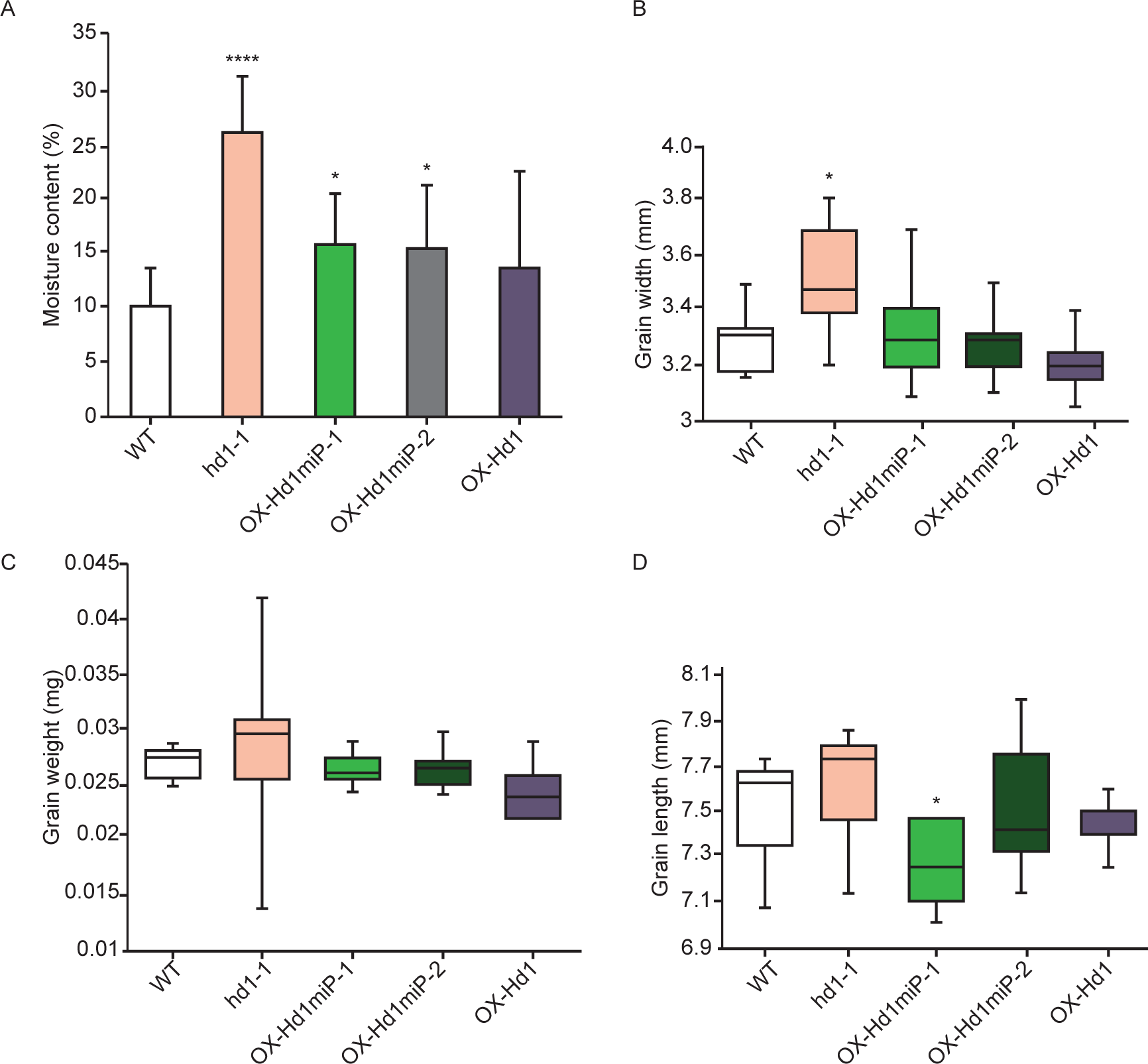
Yield analysis of Hd1 overexpressors. (A) Moisture content of WT, hd1-1, OXHd1miP, OX-Hd1. (B) Grain width of WT, hd1-1, OX-Hd1miP, OX-Hd1. (C) Grain weight of WT, hd1-1, OX-Hd1miP, OX-Hd1. (D) Grain length of WT, hd1-1, OX-Hd1miP, OX-Hd1.

Comparison of the weight, width and length of the harvested grains showed that all the lines produced grains with similar width and length with the exception of *hd1-1* knockout plants which featured a slightly wider grain compared to WT (Figure 4b, 4c and 4d). Since *hd1-1* grains also showed higher moisture content than the other grains, the larger grain size could be attribute to the higher moisture content. Taken together these results shows that a comparison of *Hd1miP* to WT, overexpressors and knockouts reveals yield traits that may be mediated by Hd1 on a protein level.

## CONCLUSION

Synthetic miPs are proving to be a useful tool in targeting and inhibiting proteins in a more specific manner with a reduced likelihood of an off-target effect. Using miP as a regulatory tool involves knowing the interaction domain of the protein of interest and the ability of the miP to form inactive heterodimers with the target; this is further complicated if there are other protein components or processes that are affected by the target. Synthetic miPs can also aid in deciphering what processes are dependent on the activity of the inhibited target protein. As is observed in OX-*Hd1miP*, miPs as a regulatory tool can be extended to crop plants the phenotypes of which can be observed in pathways that are involved.

## MATERIALS AND METHODS

### Generation of Transgenic Plants

To generate *OX-Hd1* plants, the Hd1 coding sequence was amplified from leaf cDNAs using primers listed in Supplemental Table 1 and cloned into Invitrogen vector pDONR201. The coding sequence was subsequently recombined into plant expression vectors pH2GW7 plasmid under the control of the constitutive rice ACTIN2 promoter. The *OX-Hd1miP-1* plants were generated by the amplification of a truncated form of Hd1 using primers that amplify the cDNA from the start to 429bp downstream of the start codon. This was fused to the cDNA coding for 3XFLAG by PCR, cloned into pDONR201 and recombined into pH2GW7 (LR recombinase mix, Invitrogen).

The clones were transformed into Nipponbare embryonic calli produced seeds using the EHA105 strain of Agrobacterium tumefaciens according to the protocol of Sahoo et al. (Sahoo, Tripathi, Pareek, Sopory, & Singla-Pareek, 2011) Transgenic plants were selected on 50 mg/L hygromycin during selection I and 100 mg/L hygromycin during selections II.

### Growth Conditions and Sampling

All rice (*O*. *sativa*) lines *OX-Hd1*, *OX-Hd1miP*, *hd1-1* and WT were grown under SD (10L/14D), LD (14.5L/9.5D) and SLD 16.5L/7.5D) in reach in climate chamber. Heading date was scored at panicle emergence. The plants were grown at a temperature of 28° and humidity of 80% during the day and temperature of 24°with 90% humidity at night.

### Protein expression and protein interaction assays

For protein expression analysis of Hd1 in *OX-Hd1* plants, plant material was harvested from 4-week-old *OX-Hd1* plants, flash frozen with liquid N2 and crushed with steel beads. Crushed material was dissolved is protein extraction buffer (50mM Tris HCL (pH7.5), 0.1% SDS, 150mM NaCl, 2mM EDTA, 2% BME, 4M urea, and protease inhibitor (cOmpleteTM, EDTA-free Protease Inhibitor Cocktail tablet). The supernatant was obtained for western blot analysis after 15 mins incubation on ice and centrifugation. For expression analysis of Hd1miP in *OX-Hd1miP* plants. Plant material plant was harvested from 4-weeks-old OX-*Hd1miP* plants, frozen with liquid N2 and crushed by metal beads. Crushed material was dissolved is protein extraction buffer and pulled down using M2- antiFLAG magnetic beads (Sigma). Protein bound beads were washed with extraction buffer lacking urea and inhibitors and eluted wit. For western blot analysis mouse-anti-FLAG (1:5000) and Goat anti-mouse (1:5000) were used as primary and secondary antibodies respectively. Luminescence was detected using Super signal west pico chemuluminescent substrate

For interaction studies of Hd1 and Hd1miP proteins the interaction was tested using the Matchmaker Gold yeast-two-hybrid system (Clontech). The coding sequence of either Hd1 or Hd1miP in pDONR201 was recombined into the *pGADT7-GW/pGBKT7-GW* (LR recombinase mix, Invitrogen) using the Matchmaker yeast two-hybrid system 452 (Clontech). Bait constructs were transformed into the yeast strain YM4271 MATa and preys into the yeast strain PYJ654 MATα using the lithium acetate method (Gietz & Schiestl, 1991). Mating and growth of diploid yeast was performed and screened on drop out media lacking His, Leu, and Trp with concentrations of 0, 5 and 10 mM 3-aminotriazole (3-AT).

### Grain yield analysis

All materials harvested for yield analysis was harvested from plant lines (*OX-Hd1*, *OX-Hd1miP*, *hd1-1* and WT) grown in LD (14.5L/9.5D) conditions. All Panicles in the plant was harvested from each plant 35days after first panicle emergence in the plant. Approximately 8 replicates was measured in each line. To measure the number of panicles all the panicles per plant was counted. To measure fertility rate all seeds were harvested from each panicle and number of filled seeds in all panicles per plant was used to determine the to the fertility rate. The number of filled seeds per mature panicle was used to determine fertility rate of mature panicles. For measuring moisture content, the weight of 10 seeds/plant was measured at harvest then dried at 60 degrees for 4 hours and weighed again. This represented the starting normalized weight. The seeds were subjected to another drying overnight at 105 degrees (bone dry) then weighed. For measuring the width and length of the grains 10 seed per line was arranged on graphing paper and their width and length was measured.

---

**Table 1**: Measurements of number of mature grains, panicles, and spikelet fertility WT, hd1-1, OX-Hd1miP, OX-Hd1.

